# The SMAC Mimetic AZD5582 is a Potent HIV Latency Reversing Agent

**DOI:** 10.1101/312447

**Authors:** Gavin C. Sampey, David M. Irlbeck, Edward P. Browne, Matthew Kanke, Alecia B. McAllister, Robert G. Ferris, Jessica H. Brehm, David Favre, Jean-Pierre Routy, Corbin D. Jones, Nancie M. Archin, David M. Margolis, Richard M. Dunham

**Author notes:** Corresponding author; Genetic Medicine Bldg., CB 7042, 120 Mason Farm Rd, Rm 2096, Chapel Hill, NC 27599-7042. **Conflicts of Interest** David M. Irlbeck, Robert G. Ferris, Jessica H. Brehm, David Favre, and Richard M. Dunham are employees of GlaxoSmithKline. All other authors declared that no conflict of interest exists.

## Abstract

The leading strategy towards eradication of human immunodeficiency virus (HIV) infection is the depletion of viral reservoirs through reversal of viral latency, followed by clearance of persistently infected cells. To date, a latency reversing agent (LRA) that reactivates a majority of the quiescent provirus population, without significant off-target effects, has not been identified. We show here that molecules mimicking the active N-terminal tetrapeptide of the second mitochondrial-derived activator of caspases (SMACm) potently reverse HIV latency *in vitro* and *ex vivo* without the pleotropic cellular effects seen with other LRA. We verified that SMACm facilitate latency reversal through activation of the non-canonical NFκB pathway as exemplified by rapid degradation of cellular inhibitor of apoptosis protein 1 (cIAP1), followed by a slower conversion of the inactive p100 form of NFκB2 into the active p52 transcription factor. A potent representative of this class, AZD5582, increases cell-associated HIV RNA expression in resting CD4+ T cells from ART-suppressed, HIV-infected donors while altering the expression of a restricted number of human genes. These findings represent the first demonstration that SMACm have single agent latency reversal activity in patient-derived cells and support evaluation of this class of molecules in preclinical animal models.

## Introduction

While antiretroviral therapy (ART) has greatly reduced the mortality and morbidity in HIV-infected patients, a functional or sterilizing cure remains elusive. A persistent reservoir of latently infected cells remains during ART, and viremia usually returns within a few weeks following cessation of ART (1). This reservoir decays slowly, with a half-life of 43 to 44 months (2, 3), making viral eradication unlikely despite decades of ART. As such, strategies to accelerate decay of the reservoir are under intense investigation. One such strategy aims to reverse HIV latency, induce proviral RNA and protein expression, and render the formerly latently infected cells vulnerable to viral cytopathic effects or clearance by the immune system (4). To date, concerns that current latency reversing agents (LRA) tested *in vivo* may fail to act on the majority of quiescent HIV proviruses or may do so at the cost of significant off-target effects have constrained testing of LRA *in vivo*.

LRA studied to date rely on either broad activation of cellular pathways such as NFκB, MAPK, and calcineurin (5, 6), or modification of repressive epigenetic marks on chromatin around the latent provirus (7-11). However, these mechanisms have pleiotropic cellular and systemic effects, and lack selectivity for reactivation of HIV. Recently, the targeting of the non-canonical NFκB (ncNFκB) pathway to reverse latency in cell line models was reported (12). The ncNFκB pathway is typically activated by ligation of a subset of TNF receptor family members (13). In the steady state, a multimolecular complex with ubiquitin ligase activity consisting of TNF receptor-associated factor 2 (TRAF2), TRAF3, and cellular inhibitor of apoptosis protein-1 (cIAP1) associates with the cytoplasmic portion of the unligated receptor and constitutively ubiquitinylates and degrades the NFκB-inducing kinase (NIK). Upon receptor ligation, cIAP1 ubiquitinylates TRAF3 and auto-ubiquitinylates, leading to proteasomal degradation of TRAF3 and cIAP1, thereby disinhibiting NIK accumulation. NIK is constitutively active and, once accumulated, phosphorylates the inhibitor of κB kinase-α (IKKα) homodimer. The activated IKKα/IKKα homodimer then phosphorylates the inactive p100 form of NFκB2 leading to ubiquitinylation by Skp1-Cul1-F-box ubiquitin ligase (SCF^βTrCP^) and proteasomal cleavage of p100, releasing the active p52 subunit. p52 associates with RelB, and this heterodimer translocates into the nucleus to drive transcription from κB promoter elements.

In addition to receptor ligation, ncNFκB can be activated by signaling intermediates of the apoptosis cascade (14, 15). Cleavage of the second mitochondrial activator of caspases (SMAC) from the mitochondrial membrane exposes the N-terminal motif Ala-Val-Pro-Ile, which binds specifically to the baculovirus intermediate repeat (BIR) domains of the IAP proteins. Such BIR binding in cIAP1/2 activates the ubiquitin ligase activity of the TRAF2:TRAF3:cIAP complex, inducing autoubiquitinylation and degradation of cIAP1/2 (16), NIK accumulation, and activation of the ncNFκB pathway. Binding of SMAC to the BIR domains of XIAP and ML-IAP antagonizes the caspase inhibition activities of these molecules, often overexpressed in tumor cells, leading to potentiation of apoptosis (17-19). As such the Ala-Val-Pro-Ile motif of SMAC has been the subject of significant attention in oncology, leading to discovery of a class of peptide mimetics that have SMAC-like activity, referred to as SMAC mimetics (SMACm). SMACm potently activate the ncNFκB pathway and do not induce apoptosis in non-tumor cells, and as such are of interest to reverse HIV latency (12).

Here we demonstrate that the SMACm AZD5582 functions alone as a potent LRA *in vitro* and *ex vivo* while, in contrast to other well-studied LRA, having limited additional effects on CD4+ T cells. Using a Jurkat reporter cell line model of latency, we show that several members of a panel of commercially available SMACm reverse HIV latency. Potent SMACm were confirmed to engage cIAP1 and activate the ncNFκB pathway in primary CD4+T cells. Furthermore, AZD5582 treatment had minimal impact on global gene expression in isolated CD4+ T cells. Exposure of total and resting CD4+T cells from ART-suppressed, HIV infected subjects to AZD5582 led to a significant increase in cell-associated full-length HIV RNA (caRNA). Moreover, *ex vivo* AZD5582 exposure at durations and concentrations achievable *in vivo* led to engagement of cIAP1, activation of ncNFκB, and induction of caRNA. In sum, our work demonstrates that SMACm have HIV latency reversal activity with minimal off-target effects and therefore should be evaluated in preclinical models as part of an HIV cure strategy.

## Results

### ncNFκB activation by SMACm and minimal NIK reverses HIV latency *in vitro*

The potential for SMACm to reverse HIV latency via the ncNFκB pathway was first suggested in studies of cell-line models of latency (12). We sought to validate and extend these findings, and to this end characterized the HIV latency reversing activity of a panel of commercially available SMACm. We first tested these molecules in an assay using a pool of three selected clones of Jurkat cells that demonstrated quiescent but inducible viral expression following infection with an HIV NL4-3 engineered with a bicistronic reporter consisting of luciferase and murine heat shock antigen (mHSA) in place of nef (Irlbeck, manuscript in preparation). Dimeric SMACm such as BV6, birinapant, SM-164, and AZD5582 demonstrated the strongest combination of luciferase induction in terms of maximum level (Vmax) and potency (pEC50) amongst the panel of SMACm tested, greatly exceeding the reporter induction of comparable monomeric compounds AT406, GDC-0152, and LCL161 (Fig.1A and Table 1). Examination of the kinetics of reporter expression demonstrated that SMACm-mediated induction of luciferase was slower and less strong than that observed with canonical NFκB activating agents (Fig.1B). For example the PKC agonist GSK445, a stabilized ingenol B derivative (manuscript in preparation and ref. (20)), induced luciferase activity that peaked approximately 24 hours after exposure at 35-fold higher than DMSO alone and quickly waned, while AZD5582-induced activity plateaued at 24 hours after exposure at approximately 15-fold higher than DMSO and was sustained near that level for up to 72 hours (Fig.1B). To confirm that HIV latency reversal could be achieved through the activation of the ncNFκB pathway without SMACm, we infected a single clone of the latent HIV-infected Jurkat reporter cell described above with a lentiviral vector encoding eGFP and minimal NIK (mNIK). This mNIK construct lacks the N-terminal TRAF3 binding domain of NIK responsible for interaction with the TRAF3:TRAF2:cIAP1 complex and ubiquitin mediated degradation of NIK. In turn, mNIK accumulates and, as NIK is constitutively active, ncNFκB activation occurs (21, 22). Infection with the vector encoding both eGFP and mNIK led to spontaneous HIV LTR-driven murine HSA reporter expression among the GFP+ cells as compared to uninfected cells (GFP-) in the same culture or to cells infected with a control vector lacking mNIK (Fig.1C). Moreover, combining mNIK infection with exposure to the bromodomain inhibitor, iBET151, led to synergistic reporter activation in the GFP+ cells, but not GFP-cells (Fig.1C).

**Fig. 1.**
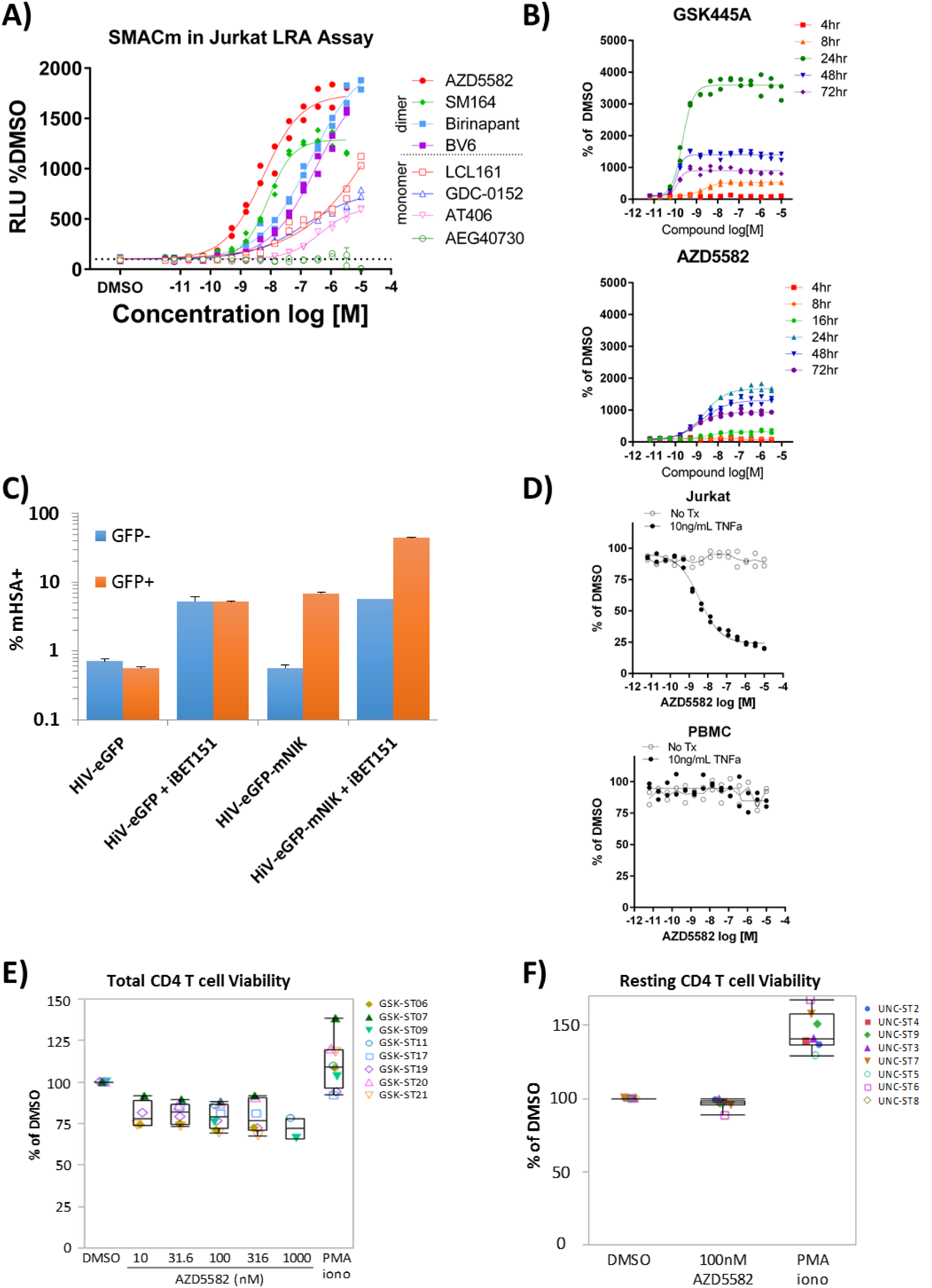
Activation of ncNFκB by SMACm and mNIK reverses HIV-1 latency in a Jurkat reporter model with minimal impact on cell viability in primary cells. (A) DMSO-normalized reporter signal induced by a dose titration of a panel of mono and bivalent SMACm Jurkat luciferase reporter model of HIV-1 latency. (B) Time course of DMSO-normalized reporter signal induced by GSK445A or AZD5582. (C) mHSA reporter expression in Jurkat cells transfected with a truncated NIK construct (HIV-eGFP-mNIK) or empty vector (HIV-eGFP), gated on cells successfully transfected (GFP+) or not (GFP-) and treated with ĨBET151 as indicated. (D) Viability in Jurkat cells and primary PBMCs treated with AZD5582 in the presence or absence of 10ng/mL TNFα. (E) DMSO-normalized viability as measured by Cell Titer Glo assay from total CD4+ T cells isolated from 10 patients across all five AZD5582 concentrations tested. (F) DMSO-normalized viability as measured by Cell Titer Glo assay from resting CD4+ T cells isolated from eight patients and treated with 100nM AZD5582.

**Table 1.**
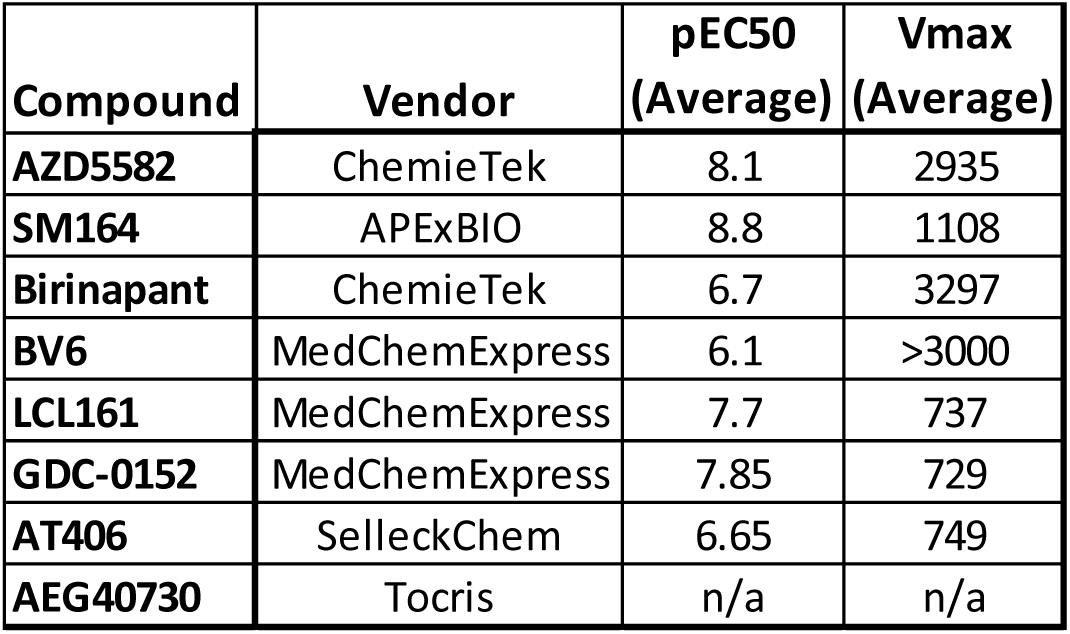
**Summary of SMACm Activities in the Jurkat Luciferase Assay**

As SMACm were initially developed to potentiate apoptosis, we studied the impact of the most potent SMACm tested, AZD5582, on cellular viability in cell lines and primary cells. Using a luciferase-based assay for quantifying ATP content, we found that AZD5582 did not significantly impact viability of Jurkat cells. However, addition of a sub-cytotoxic concentration of TNFα (10ng/mL) led to AZD5582 dose-dependent cell death in Jurkat cells, with a CC50 comparable to that observed for AZD5582 LRA activity in the Jurkat reporter cells (Fig.1D), as described for SMACm and TNFα exposure in other transformed cell lines (23). AZD5582 had minimal impact on cellular viability in PBMC and addition of TNFα did not potentiate cell death, in contrast to what was observed for Jurkat cells (Fig.1D). Similar to total PBMC, SMACm did not lead to overt cell death among isolated total CD4+ T cells (Fig.1E) or resting CD4+ T cells (Fig.1F), though a consistent decrease in ATP levels (approximately 80% of DMSO treated cells) was observed among total CD4+ T cells (Fig.1E). Together these findings confirm earlier reports that SMACm can reverse HIV latency in cell line models and do not have a significant impact on viability in primary cells.

### Kinetics and dose-dependent activation of the ncNFκB pathway by SMACm

To confirm engagement of the ncNFκB pathway by SMACm, we assessed depletion of the proximal SMACm target proteins – cIAP1 and cIAP2 – and downstream processing of p100 to p52 in primary CD4+ T cells exposed to AZD5582. Immunoblot analysis showed rapid and durable depletion of cIAP1, and to a lesser extent cIAP2, within 30 minutes of exposure to 100nM AZD5582 (Fig.2A). Although cIAP1 and cIAP2 depletion occurred rapidly, processing of p100 to p52 was not observed until 4 hours post-exposure and accumulated over 48 hours of exposure, with final p52 levels greater than 10-fold higher than observed before treatment (Fig.2B). The canonical NFκB pathway was not activated by AZD5582, as evidenced by maintenance of IκBα, and instead IκBα protein levels increased over time during AZD5582 exposure (Fig.2B). The depletion of cIAP1 and conversion of p100 to p52 by AZD5582 was dose dependent, with cIAP1 depleted and p100 cleaved at concentrations exceeding 0.1nM. Despite dose dependent depletion of cIAP1 in primary CD4+ T cells comparable to that observed with AZD5582, cleavage of p100 into the active p52 was only observed at birinapant exposures greater than 100nM (Fig.2D-E). The full activation of ncNFκB, as measured by p100 to p52 conversion, by AZD5582 at comparable concentrations to those where HIV latency is reversed supports the conclusion that the latency reversal activity is mediated through ncNFκB activation.

**Fig. 2.**
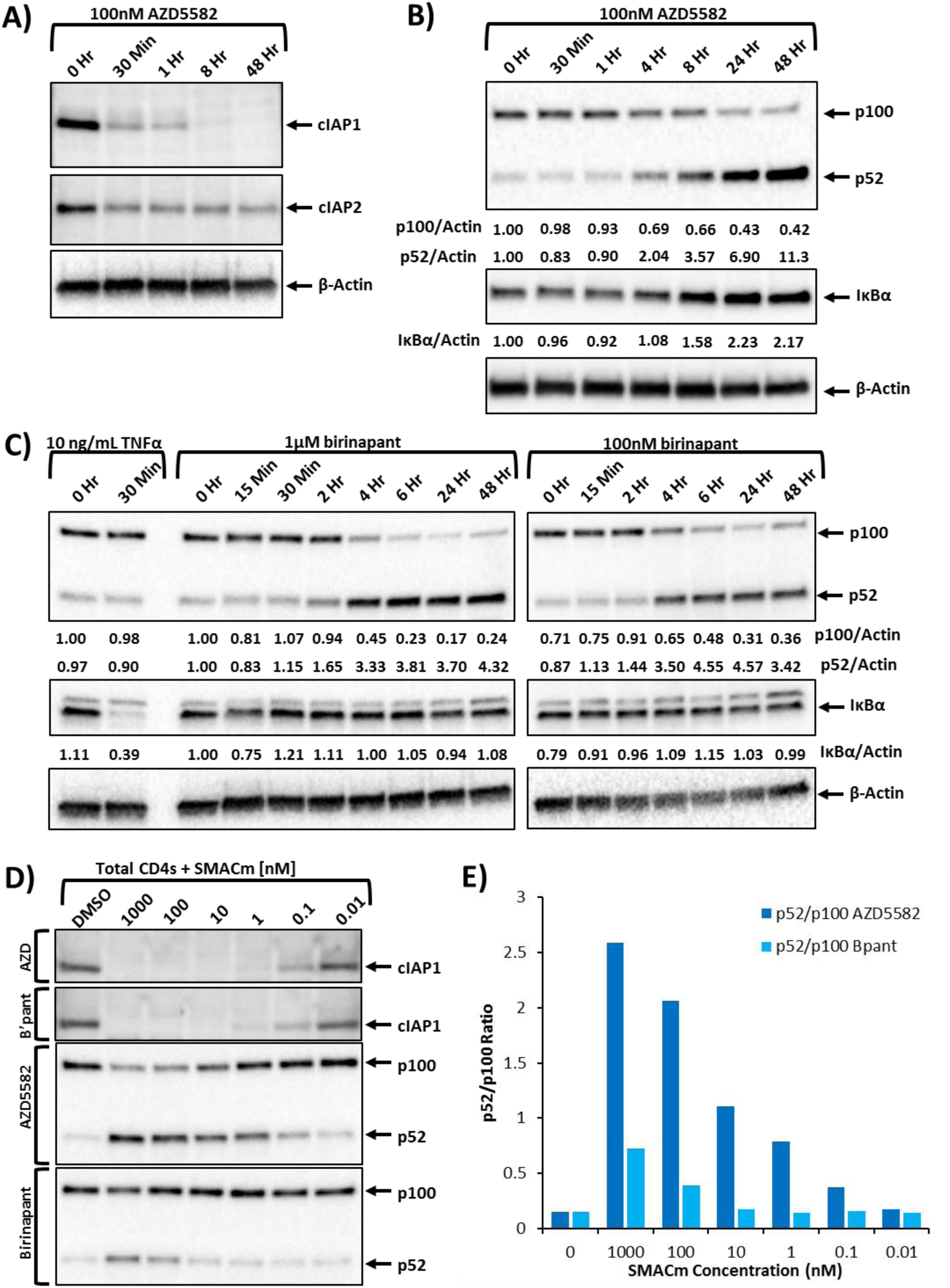
SMACm rapidly degrade cIAP1 with slower kinetic conversion of inactive p100 into active p52. (A-B) Immunoblot analysis of isolated total CD4+ T cells treated with 100nM AZD5582 examining components of the canonical and ncNFκB pathway over a 48-hour time course post-treatment. (C) Immunoblot analysis of canonical and ncNFκB components in Jurkat cells over time after treatment with TNFα or 1μM or 100nM birinapant. (D) Total CD4+ T cells were treated with a broad range of concentrations (10pM to 1μM) of the SMACm AZD5582 or birinapant, and cell lysates were analyzed by immunoblot, probing for ncNFκB components. (E) The ratio of the p52 to p100 band densities shows the higher potency of AZD5582 over birinapant in activating the ncNFκB pathway.

### Short duration exposure to AZD5582 activates the ncNFκB pathway and reverses HIV latency

AZD5582 is cleared from circulation rapidly; relevant *in vivo* doses are estimated to achieve only one to two hours of plasma exposure at concentrations greater than the EC_50_ (24). To study whether such brief exposure supports latency reversal, we treated cells with AZD5582 for short periods of time followed by extensive washing to remove the drug and measured ncNFκB pathway activation over time by western blot. In primary CD4+ T cells, as short as one hour exposure to 100nM AZD5582 was sufficient to deplete cIAP1 and cIAP2 and induce processing of p100, with the densitometric ratio of p52 to p100 in western blot achieving greater than 60% of that observed with continuous exposure (Fig.3A-E). Similarly, exposure of the Jurkat reporter cells for as little as 30 minutes to AZD5582 led to induction of luciferase activity comparable with that observed with continuous AZD5582 exposure (Fig.3F). These pulse-wash studies demonstrate that exposure to SMACm at intervals and concentrations achievable *in vivo* can activate ncNFκB and reverse HIV latency.

**Fig. 3.**
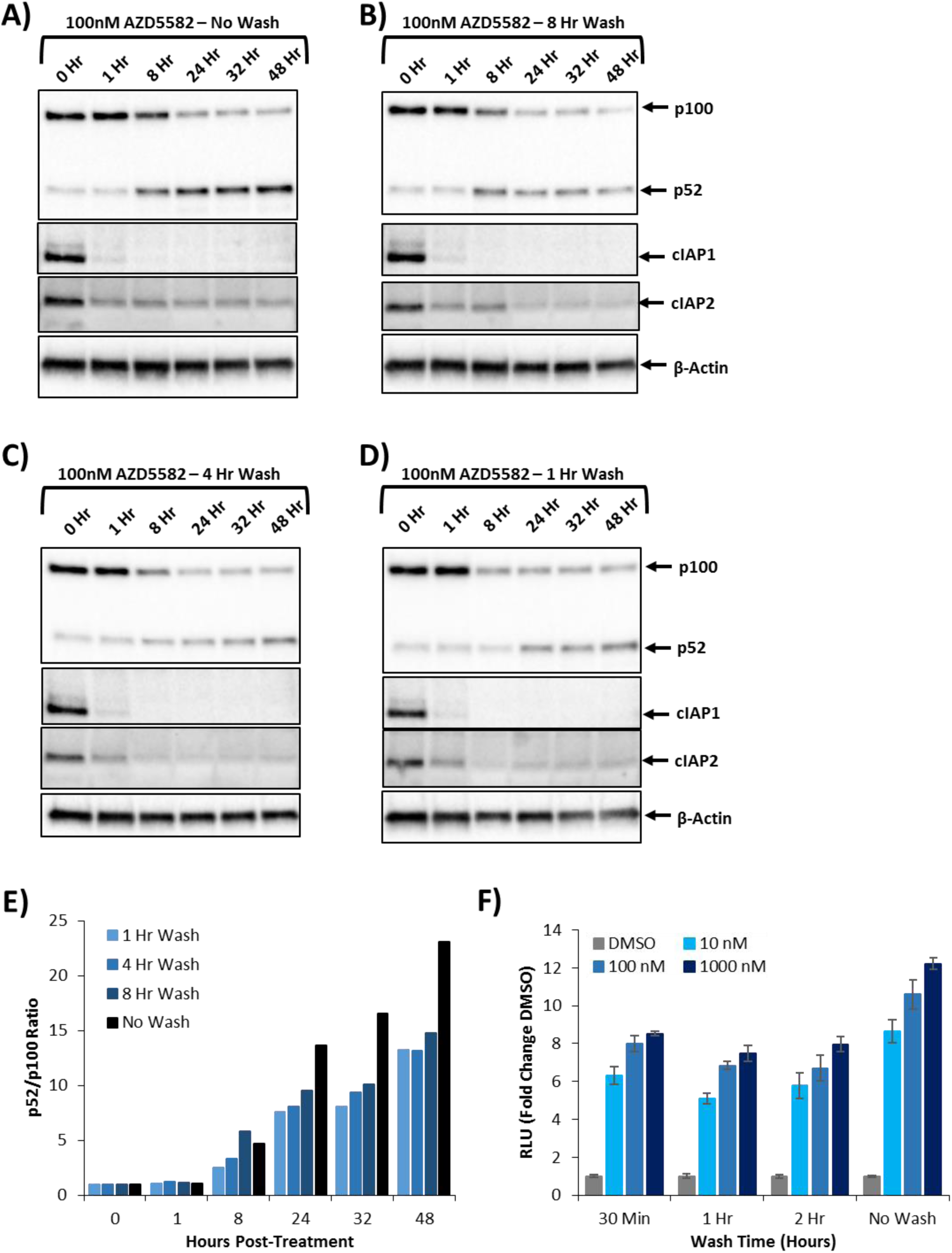
Immunoblot analysis of total CD4+ T cells and latency reversal of the Jurkat reporter after pulse-wash treatment with AZD5582. Isolated totaled CD4+ T cells treated with 100nM of AZD5582 and then either not washed (A) or washed three times with PBS at eight (B), four (C), or one hour (D) posttreatment. Whole cell lysates were then analyzed by immunoblot for components of the ncNFκB pathway. (E) Densitometry analysis of the ratio of p52 to p100 from the pulse-wash assay immunoblots. (F) DMSO normalized fold-change in luciferase activity from the Jurkat reporter model after exposure to AZD5582 (10, 100, or 1000nM) for either 30 minutes, 1 hour, 2 hour, or continued exposure.

### AZD5582 has minimal impact on activation or gene expression in CD4+ T cells

Classical latency reversal is achieved by maximal T-cell activation using agents such as anti-CD3/anti-CD28 or protein kinase C agonist (PKCa). These molecules induce strong activation signatures in T cells and initiate substantial changes in gene expression. To test whether SMACm have similar activity, we first measured CD69 expression in primary T cells exposed to AZD5582. We found no induction of CD69 in CD4+ T cells cultured for 24 hours in the continuous presence of up to 1000nM AZD5582 (Fig.4A). We then characterized the transcriptome of isolated CD4+ T cells treated with 100nM AZD5582 or 250nM the PKCa GSK445A (20) to better understand the global transcriptional impact of SMACm. We performed unbiased, stranded mRNA sequencing from total RNA extracted from total CD4+ T cells isolated from four ART-suppressed, HIV-infected donors. If, like in classical latency reversal approaches, AZD5582 treated cells evinced a strong change in gene expression, this would suggest that AZD5582 has a strong activation signature and potentially collateral effects on the cell. Instead, AZD5582 exposure led to relatively few differentially expressed genes when compared to vehicle (upregulated genes: 27 at 2hrs, 134 at 6hrs, 387 at 24hrs), while GSK445A led to differential expression of nearly 100-fold more genes (upregulated genes: 2643 at 2hrs, 2902 at 6hrs, 2303 at 24hrs) (Fig.4B). AZD5582 treatment primarily led to increases in gene expression, as compared to a relative balance of increased and decreased expression after GSK445A exposure (Fig.4B). In agreement with the slow, durable kinetics of p100 processing described above, the number of genes affected by AZD5582 increased over time, and only 15 genes activated at 2 and 6 hours were not up-regulated at 24 hours (Fig.4D). Among the genes induced by AZD5582 at any timepoint, more than half were also induced by GSK445A (Fig.4C). Gene ontology analysis of the shared 87 induced genes at 6 hours showed enrichment for NFκB-related pathways (Fig.4E), while no enrichment was identified for the remaining 47 genes induced by AZD5582 alone. This analysis suggests that HIV latency reversal by SMACm does not require full T-cell activation and occurs in the context of orders of magnitude fewer genes than those induced by T-cell activation by PKCa.

**Fig. 4.**
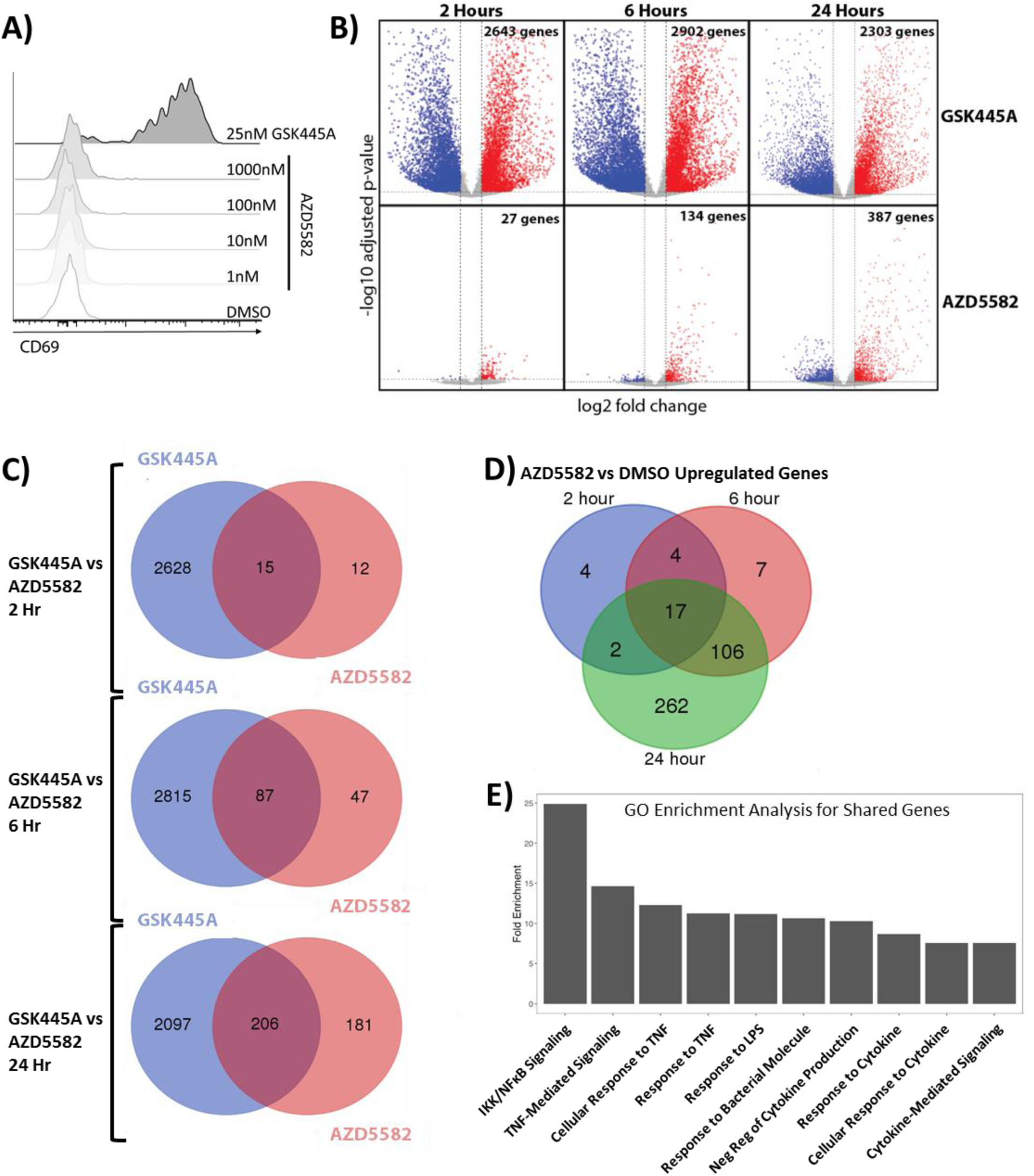
Phenotypic and transcriptomic analysis of AZD5582 and GSK445A treated total CD4+ T cells. RNA-seq analysis of total CD4+ T cells isolated from four HIV-infected ART-suppressed individuals treated ex vivo with either DMSO alone, GSK445A, or AZD5582. (B) Volcano plots showing up and down-regulated genes at 2, 6, and 24 hours post-treatment with GSK445A or AZD5582, with the log2 fold change on the x axis and logiø adjusted p-value on the y-axis (B) as compared to DMSO alone. (C) Venn diagram of the overlapping genes up-regulated by GSK445A and AZD5582 at 2, 6, and 24 hours post-treatment. (D) Venn diagram of genes up-regulated by AZD5582 at 2, 6, and 24 hours post-treatment. (E) Gene ontology enrichment analysis of up-regulated genes shared by GSK445A and AZD5582 at 6 hours post-treatment.

### AZD5582 can increase viral transcripts from patient total and resting CD4+ T cells *ex vivo*

To understand whether potent SMACm can act as LRA in patient-derived cells, we measured total full length HIV RNA levels using quantitative PCR with primers targeting gag (herein referred to as caRNA) in isolated total CD4+ T cells from ART-suppressed HIV-infected donors treated with SMACm *ex vivo* (Table 2). We observed up to a 2.2-fold induction of caRNA in at least one condition tested in six of the ten donors (Fig.5A and Supplemental Fig.1A) treated with AZD5582. Significant caRNA induction was observed in some donors with exposure to as low as 10nM AZD5582. caRNA induction was comparable between cells treated with different concentrations of AZD5582 and did not exhibit a dose response. This lack of a dose response is not unexpected, though, as all tested concentrations are above the Jurkat EC50 of 8nM. A similar study of birinapant did not lead to increased caRNA in any of the seven donors studied (Supplemental Fig.1B), as expected in the context of the observed lack of p100 cleavage in primary CD4+ T cells (Fig.2D-E). In the cells from the four donors in which caRNA was not induced by AZD5582, baseline caRNA levels were at or near the limit of quantification for the PCR assay in all conditions (7 copies per reaction), or caRNA induction was not observed following maximal PMA/Ionomycin treatment, indicating that induction may have occurred but was below the limit of detection (Supplemental Fig.1A). When the data from all ten donors was considered in aggregate, statistically significant induction was observed as low as 31.6nM (p<0.01, Fig.5A).

**Fig. 5.**
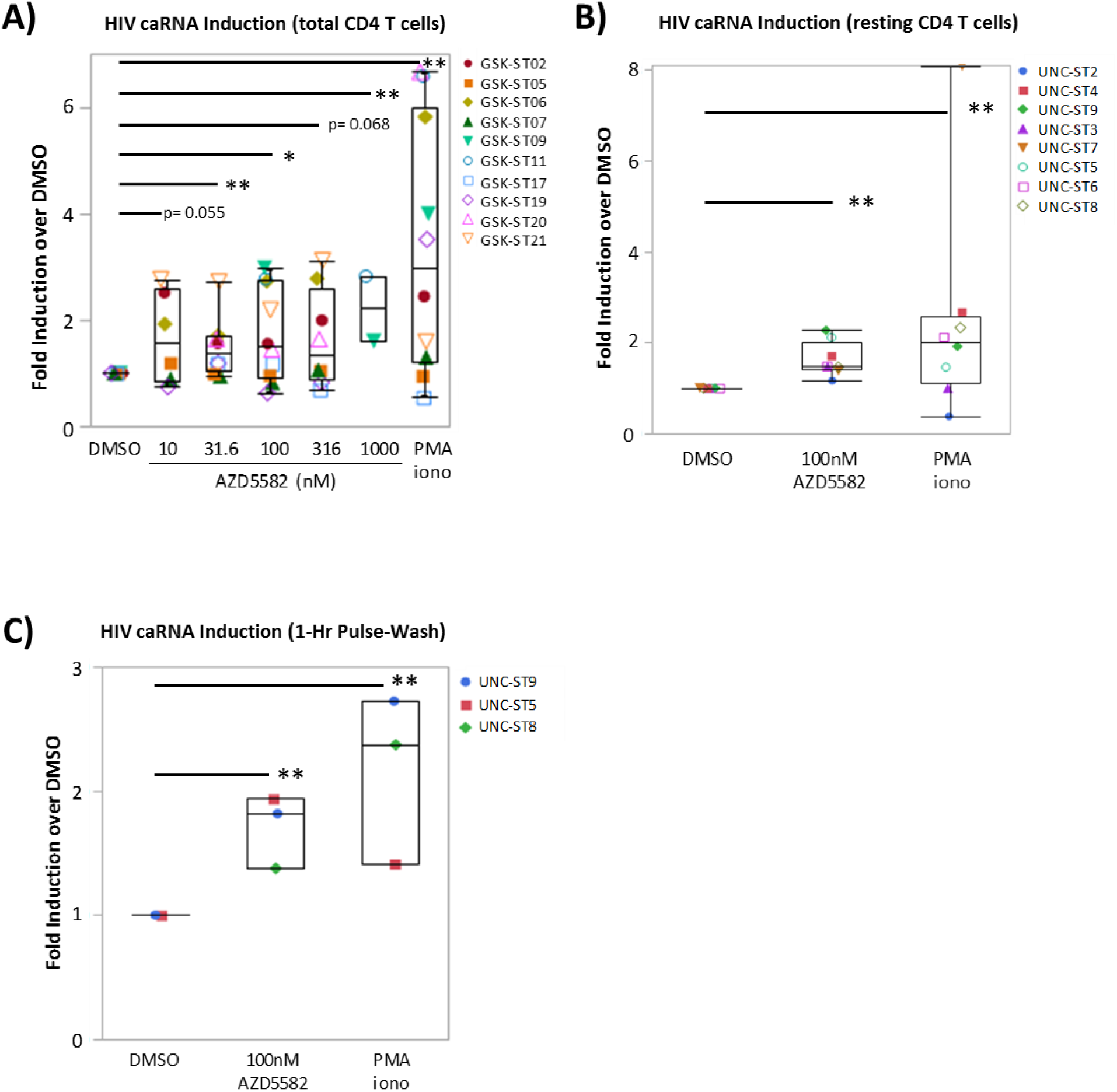
Cell associated gag RNA from total and resting CD4+ T cells treated with AZD5582. Total or resting CD4+ T cells were isolated from HIV-infected, ART-suppressed individuals and treated with AZD5582 and then ca-HIV RNA copies were quantified by RT-qPCR. (A) Combined fold-change in caRNA from total CD4+ T cells isolated from 10 patients and treated with 5 AZD5582 concentrations for 48 hours. All conditions tested in biological triplicates and each sample run in technical triplicate. The caRNA copies per million cells were normalized to the DMSO treated negative control and PMA/I was treatment was used as a positive control. (B) Fold change in caRNA copies from stably treated patient resting CD4+ T cells treated with DMSO, 100nM AZD5582, or PMA/I (cells from 8 patients tested in triplicate; normalized to DMSO). (C) caRNA copies per cell after 1-hour pulse of AZD5582 in total or resting CD4+ T cells from stably treated patients. All conditions tested in biological triplicates and each sample run in technical triplicate RT-qPCR. All statistical analysis conducted using nonparametric comparisons for each pair using the Wilcoxon method. * = p-value < 0.05; ** = p-value < 0.01

**Table 2.** **Patient Characteristics**

| **ID** | **Age** | **Gender** | **Ethnicity** | **CD4 count** | **ARV Duration** |
| --- | --- | --- | --- | --- | --- |
| **GSK-ST02** | 34 | M | Caucasian | 603 | 6yrs |
| **GSK-ST05** | 32 | M | Caucasian | 417 | 6yrs |
| **GSK-ST06** | 61 | M | Caucasian | 442 | 22yrs |
| **GSK-ST07** | 48 | M | Caucasian | 409 | 16yrs |
| **GSK-ST09** | 50 | M | Caucasian | 682 | >6mo |
| **GSK-ST10** | 63 | M | Caucasian | 896 | 13yrs |
| **GSK-ST11** | 56 | M | Caucasian | 576 | 6yrs |
| **GSK-ST12** | 38 | M | Caucasian | 812 | 6yrs |
| **GSK-ST16** | 54 | M | Caucasian | 1585 | 16yrs |
| **GSK-ST17** | 48 | M | Caucasian | 377 | 15yrs |
| **GSK-ST19** | 47 | M | Hispanic | 239 | 14yrs |
| **GSK-ST20** | 53 | M | Caucasian | 176 | 25yrs |
| **GSK-ST21** | 35 | M | Caucasian | 372 | >6mo |
| **UNC-ST1** | 53.6 | M | Caucasian | 706 | 3yrs |
| **UNC-ST2** | 50.9 | M | Caucasian | 563 | 1yr |
| **UNC-ST3** | 61.9 | F | African American | 1034 | 1.7yrs |
| **UNC-ST4** | 46.8 | M | Caucasian | 516 | 2yrs |
| **UNC-ST5** | 23.3 | M | Caucasian | 668 | 2yrs |
| **UNC-ST6** | 52.1 | M | Caucasian | 608 | 10yrs |
| **UNC-ST7** | 59.5 | M | African American | 733 | 6yrs |
| **UNC-ST8** | 37.9 | F | African American | 873 | 16yrs |
| **UNC-ST9** | 28.3 | M | Caucasian | 574 | 5yrs |

Total CD4+ T cells are more likely to contain cells that are producing HIV RNA directly *ex vivo* and stimulation may simply increase this pre-existing HIV RNA expression, complicating the interpretation of caRNA induction by putative LRA. We therefore specifically examined isolated resting CD4+ T cells, in which baseline levels of caRNA are lower and which likely contain a more resistant subpopulation of latently infected cells. Indeed, it is this population that has been shown to be stable despite years of ART (2, 3). As observed with total CD4+ T cells, 48 hours of exposure to 100nM AZD5582 increased caRNA an average 1.6-fold (p<0.0001 vs. DMSO alone) in resting CD4+ T cells from 6 of 8 donors, compared to an average 2.6-fold induction by PMA/ionomycin (p<0.0001 vs. DMSO alone; Fig.5B and Supplemental Fig.1C). Finally, to determine whether short exposures to concentrations of AZD5582 achievable *in vivo* can reverse HIV latency in primary patient-derived cells, we exposed patient-derived total or resting CD4+ T cells to 100nM AZD5582 for one hour and measured caRNA 48 hours later. In the three samples tested, one hour exposure to 100nM consistently increased caRNA to levels comparable with 48-hour continuous exposure (Fig.5C). These data represent the first demonstration that SMACm alone can reverse HIV latency *ex vivo* in resting CD4+ T cells from stably ART-suppressed donors.

## Discussion

While several LRA have shown efficacy in *in vitro* and *ex vivo* experiments, the most potent of these compounds have significant collateral effects on the host cells such as overt T-cell activation and/or broad cytotoxicity. Here we demonstrate that a SMACm, AZD5582, reverses HIV latency through the activation of the ncNFκB pathway without evidence of host cell activation and with minimal host cell transcriptional changes. Moreover, we demonstrate that AZD5582 activates the most quiescent proviral reservoir of the resting CD4+ T cells at concentrations and exposure durations that are achievable in preclinical animal models.

In our initial screen of SMACm using a Jurkat reporter model of HIV latency, we observed a range of potencies across this class of compounds ranging from completely inactive (AEG40730) to an EC50 of 7.5nM (AZD5582) (Fig.1A). The bivalent SMACm compounds demonstrated higher activity and potency than did their monovalent counterparts. Based on immunoblot studies, we found that AZD5582 potently induces degradation of cIAP1 at sub-nanomolar concentrations (Fig.2D) and induces degradation of cIAP2 to approximately 50% of its pre-treatment levels (Fig.2A), comparable to reported cell-free BIR domain binding activity (25) and our observed EC50 in the Jurkat model (Fig.1A). These findings confirm previous reports (12) and support the conclusion that SMACm activate the ncNFκB pathway to reverse HIV latency. Unlike other potent LRA that typically activate rapid and potent mechanisms, SMACm led to slow and durable activation of ncNFκB, as evidenced by conversion of p100 to active p52 starting 2 to 4 hours post-treatment and continuing for at least 48 hours post-treatment (Fig.2B-C). The activation kinetics observed in this study are in line with other reports of ncNFκB activation by SMACm and generally support a less frequent dosing regimen (e.g., once or twice weekly) as has been utilized in SMACm preclinical and clinical studies (25, 26). Moreover, the durable activation of ncNFκB and observed HIV latency reversal despite limited duration exposure to SMACm (Fig.3 and Fig.5C) further boost confidence in increased dosing intervals in future *in vivo* experiments. It will be important in the future to investigate how long ncNFκB remains activated following SMACm exposure and how such activation is resolved, as these parameters will be key in determining optimal dosing intervals in preclinical and clinical studies for HIV cure.

In contrast to the transcriptional signature of PKCa, SMACm were demonstrated here to have relatively minimal impact on T-cell activation or host transcriptome (Fig.4). Even when comparing timepoints at which the most genes were induced for each treatment (6 hours for PKCa vs. 24 hours for SMACm), the PKCa GSK445A induced 7.5-fold more genes than AZD5582 (Fig.4C). This limited modulation of host gene expression induced by AZD5582, along with the lack of induction of T-cell activation markers (Fig.4A) demonstrates that AZD5582 is more selective for induction of HIV transcriptional activity and may therefore be expected to have more limited collateral effects than other agonist LRA, such as PKCa. While these T-cell transcriptomic findings are an important first study in examining the off-target impacts of SMACm, follow-up studies on the effects of SMACm on other cell types, such as monocytes and B cells, will be important to form a more complete understanding of SMACm on and off target activities prior to preclinical and clinical studies.

One key advance shown here is that SMACm can function as a single agent to induce HIV RNA expression in *ex vivo* treatment of CD4+ T cells isolated from ART-treated, aviremic patients. We observed statistically significant increases in caRNA in total CD4+ T cells as well as the more quiescent and refractory resting CD4+ T-cell population (Fig.5A-B). Moreover, as little as a one hour exposure to 100nM AZD5582, a concentration and duration achieved in murine oncology models, led to significant increases in caRNA in patient-derived resting CD4+ T cells with fold-increases similar to that observed with continuous exposure (Fig.5C). This set of *ex vivo* HIV latency reversing experiments gives the greatest insight into the potential of SMACm in a clinical setting due to the diversity of proviruses and host genetic heterogeneity encompassed in testing patient cells.

In this study, we show that the potent SMACm AZD5582 activates ncNFκB and reverses HIV latency in cell line models of HIV latency and in patient-derived resting CD4+ T cells, while modulating expression of only a limited number of host genes. Importantly, the *ex vivo* ncNFκB activation and LRA activity are induced with AZD5582 concentrations and durations of exposure that are achievable in preclinical models. In sum, our work demonstrates that SMACm are an important novel LRA specifically activating the ncNκKB pathway that may accelerate decay of the HIV reservoir as part of an HIV cure strategy. These studies therefore set the stage for preclinical evaluation of SMACm in animal models of HIV latency.

## Methods

### Preparation of Jurkat HIV-luciferase cell clones

Cell clones Jurkat-C16 and Jurkat-I15 were made using HIV-1 engineered to express a luciferase reporter in place of the HIV-1 nef gene (NLCH-Luci). The Jurkat-N6 cell clone was made using the same virus as described above with an additional mouse heat stable antigen CD24 (HSA) reporter located just downstream of the luciferase open reading frame and separated by a T2A element (NLCH-Luci-HSA). NLCH, the parent molecular infectious clone used to make the Jurkat clones, is a modification of HIV-1 NL4-3 (GenBank U26942) where flanking sequences were removed. All viruses were derived by transfection of human embryonic kidney 293 cells (HEK 293T) with 1μg of the HIV NL4-3 derived infectious molecular plasmid DNAs using the FuGENE HD Transfection reagent (Promega, Madison, WI) per the manufacturer’s recommendations. Supernatants were collected 48 hours post transfection, passed through a 0.2micron filter, and used to infect wild-type Jurkat cells. Following infection, cells expressing high levels of HIV-encoded HSA were removed using anti-mouse CD24 antibody clone 30-F1 (Biolegend, San Diego, CA) that was adsorbed to streptavidin-labeled magnetic Dynabeads M-280 (Life Technologies, Carlsbad, CA) covalently bound to biotin-labeled rat ant-mouse CD24 clone M1/69 antibody (BD Biosciences, San Jose, CA) per the manufacturer’s recommendations. Negatively selected, HIV-infected Jurkat cells were then limit-diluted at 0.5 cells/well in 96-well plates, and individual cell clones were expanded for two to four weeks in culture in the presence of 500nM efavirenz (EFV).

### Cell Culture and Jurkat HIV-Luciferase Assay

Jurkat HIV-luciferase clones were maintained in RPMI medium 1640 (Gibco by Life Technologies, Grand Island, NY) containing 10% (vol/vol) fetal bovine serum (SAFC/Sigma-Aldrich, Buchs, Switzerland) and 25 units/mL penicillin, 25 units/mL streptomycin (Gibco by Life Technologies, Grand Island, NY), and were split 1:4 every 3 to 4 days to maintain a cell density of ~0.3 to 1 million cells/mL. The Jurkat clones were maintained with the addition of 500nM EFV in the medium. Three Jurkat cell clones (C16, I15, and N6), each harboring one or two integrated HIV proviruses expressing the luciferase reporter gene, were added at equal amounts for a total of 5,000 cells per well to 384-well plates containing compound titrations. Dose-response testing was performed on compounds dissolved in dimethyl sulfoxide (DMSO; Fisher Scientific, Merelbeke, Belgium) dispensed in duplicate serial 3-fold, 15-point titrations using a D300e Digital Droplet Dispenser (Hewlett-Packard, Singapore) to give final assay concentrations of 10 μM to 2.1 pM in 50 μL of medium at 0.5% DMSO (vol/vol) final concentration. Cells and compound were incubated at 37°C for 48 hours followed by the addition of 20 μL of Steady-Glo^®^ Luciferase (Promega, Madison, WI). Luminescence resulting from the induction of the virally expressed luciferase was measured using an EnVision 2102 Multilabel Plate Reader (Perkin Elmer, Shelton, CT). Dose-response relationships were analyzed with GraphPad PRISM 6 using a four-parameter model to calculate the concentration of compound that gives half-maximal response (EC_50_) and the maximal percent activation compared to the vehicle control (V_max_).

### Minimal NIK Construction

Coding sequence for amino acids 330-679 corresponding to the catalytic core of NIK (mNIK) was cloned into the pHIV-eGFP vector (Addgene), a self-inactivating lentiviral expression vector that was originally generated by the Welm lab (University of Utah). This vector contains an internal EF-1a promoter that drives expression of heterologous inserts, as well as eGFP from an IRES element. To generate virus from this plasmid, pHIV-eGFP-mNIK or empty pHIV-eGFP was transfected into HEK 293T cells using Mirus LT-1 reagent (Mirus Bio), along with packaging plasmids PAX2 and MD2-VSVG. Two days after transfection, supernatant was harvested from the HEK 293 cells and clarified by low speed centrifugation for 5 minutes, followed by filtration through a 0.45μM filter (Pall Corp). Jurkat N6 cells were then plated in 12-well plates at 0.5 million cells/mL and infected with 1mL of virus supernatant for 2 hours. After 2 hours, cells were washed and cultured in RPMI with 10% FCS. At 4 days post infection, a portion of the cells were stimulated with the bromodomain inhibitor iBET151 at 10μM or DMSO for 24 hours. Reactivation of the integrated HIV reporter construct was then assessed by staining for murine HSA with an anti-HSA-PE antibody (Biolegend) at 1:100 for 30 minutes. The cells were then washed in PBS, fixed in 4% PFA, and analyzed on a LSRFortessa (Beckton Dickson) flow cytometer.

### Cell Associated HIV RNA

Peripheral blood mononuclear cells (PBMCs) were isolated from leukocytes obtained by continuous-flow leukapheresis. Total CD4+ T cells were isolated from frozen PBMCs by negative selection using the EasySep Human CD4+ T cell Enrichment kit (StemCell, Vancouver, Canada) per the manufacturer’s recommendations. Resting CD4+ T cells were isolated by negative selection with an immunomagnetic column as described previously (27) and either cryopreserved immediately or maintained for two days in IMDM medium (Gibco), 10% FBS, 2μg/mL IL-2 (Peprotech), and antitretrovirals to prevent viral expansion. For each LRA treatment, three biological replicates of 5×10^6^ total or resting CD4+ T cells were treated for 48 hours at 37°C in 1mL of RPMI medium 1640, 10% FBS, 10μg/mL of enfuvirtide (Sigma), and 200 nM rilpivirine. 10nM phorbol 12-myristate 13-acetate (PMA; Sigma) with 1μM ionomycin (Sigma) was used as a positive control for LRA activation and 0.2% DMSO vehicle was used as a negative control. Following treatment, cells were lysed and RNA and DNA were co-extracted using an AllPrep 96 RNA/DNA kit (Qiagen, Valencia, CA) per the manufacturer’s instructions, adjusting the volume of lysis buffer to 0.6mL, adding an on-column DNase I treatment (Qiagen) and eluting RNA in 50μL of water.

RT-qPCR was performed in triplicate for each of three biological replicates using TaqMan Fast Virus 1-step RT-qPCR Master Mix (Applied Biosciences) with 5μL isolated RNA and 900nM of HIV capsid primers HIV-gag (5’-ATCAAGCAGCTATGCAAATGTT-3’) and gag reverse (5’-CTGAAGGGTACTAGTAGTTCCTGCTATGTC-3’) and 250nM of FAM/ZEN/IABFQ HIV gag probe (5’-ACCATCAATGAGGAAGCTGCAGAATGGGA-3’). Samples were amplified and data was collected using a QuantStudio™ 3 Real-Time PCR system (Applied Biosystems) with the following cycling conditions: one cycle at 50°C for 5 minutes (reverse transcription), one cycle at 95°C for 20 seconds (reverse transcriptase inactivation), and 50 cycles at 95°C for 3 seconds and 60°C for 20 seconds (denaturation and annealing/extension).

HIV absolute HIV gag RNA copies per reaction were determined using an HIV gag gBlock qPCR standard (5’-ATCAAGCAGCCATGCAAATGTTAAAAGAGACCATCAATGAGGAAGCTGCAGAATGGGATAGATTGCATCCAGTGCATGCAGGGCCTATTGCACCAGGCCAGATGAGAGAACCAAGGGGAAGTGACATAGCAGGAACTACTAGTACCCTT CAG-3’ Integrated DNA Technologies), and copies were normalized to cell counts as determined by bright field microscopy. RT-qPCR efficiency was required to be between 90% to 110%. Assay values with a positive signal that were less than the lower limit of detection (LLOD) of seven HIV gag copies per reaction were adjusted to the LLOD. Analysis was performed using QuantStudio™ Design and Analysis Software (Applied Biosystems) and JMP 12.2 Statistical Discovery™ software (SAS).

### Immunoblot Analysis

For the immunoblot assays, 10 μg of cell lysate was loaded per well into 4-20% Tris-Glycine SDS-PAGE gels. Protein from the SDS-PAGE gels were transferred to Turbo Midi PVDF Transfer Packs (BioRad) using the “Mixed MW” protocol for one Midi Format Gel (constant 2.5A up to 25V, for 7 minutes) of the Trans-Blot Turbo Transfer System (Bio-Rad) with premade Trans-Blot per the manufacturer’s instructions. After transfer, PVDF membranes were blocked in 5% bovine serum albumin (BSA) in 1x TBS (BioRad) with 0.1% Tween-20 for 1 hour at room temperature with gentle rocking. Primary antibodies were added and incubated overnight at 4°C (anti-cIAP1, Abcam #ab108361, 1:1000; anti-p100/p52, Cell Signaling Technology #3017, 1:1000; anti-IκBα, Cell Signaling Technology #4812, 1:1000; anti-cIAP2, Abcam #ab32059, 1:1000; anti-Actin-HRP conjugate, Abcam #49900, 1:30,000). Following primary staining, the membrane washed three times with 1x Tris Buffered Saline (TBS)+0.1% TWEEN^®^ 20 for 10 minutes each wash. After washing, the membrane was incubated in 5% BSA in 1x TBS+0.1% TWEEN^®^ 20 with the appropriate secondary antibody for 2 hours at room temperature. Following secondary stain the membrane was washed twice for 10 minutes with 1x TBS+0.1% TWEEN^®^ 20 followed by a 10-minute wash with 1x TBS. The membrane was then patted dry with filter paper and an image was captured of the undeveloped membrane on the ChemiDoc MP Imaging System using Image Lab software (BioRad). Sufficient ECL reagent (GE Healthcare) was used to cover the membrane and a series of images were taken starting with 0.001 second and doubling to tripling the exposure time until the luminescence from the developed membrane saturated the image. The developed membrane was then washed three times with 1x TBS for 5 minutes to remove the residual ECL reagent and then stored at 4°C in sufficient 1x TBS to submerse the entire membrane. Densitometry of images of the developed membrane were then carried out using the Image Lab software. Some membranes were stripped for one minute with One Minute Plus Western Blot Stripping Buffer (GM Biosciences) and then washed three times for 10 minutes with 1x TBS. The stripped membranes were then blocked in 5% BSA in 1x TBS+0.1% TWEEN^®^ 20 for an hour and reprobed overnight with a new primary antibody.

### Flow Cytometry

Cryopreserved PBMC were thawed, plated, and exposed to a dose range of AZD5582 or GSK445A. After 24 or 48 hours of culture, cells were washed twice and stained with a cocktail of fluorophore-conjugated antibodies to 30 minutes at 4°C (anti-CD3 SP34-2 Alexa 700, BD Biosciences, 1:50; anti-CD4 L200 BrilliantViolet 711, BD Biosciences, 1:50; anti-CD8 RPA-T8 BrilliantViolet 786, 1:50; anti-CD69 FN50 BrilliantViolet 421, BD Biosciences, 1:50) and LIVE/DEAD Fixable Aqua Dead Cell Stain (ThermoFisher). Cells were washed twice and then fixed with 0.5% paraformaldehyde in PBS. Acquisition was performed on an LSRFortessa (Becton Dickinson) and analysis was carried out with FlowJo (v9.7.3).

### RNA-seq

Total CD4+ T cells were isolated from the PBMCs of four ART-treated, aviremic patients (patient IDs: GSK-ST06, GSK-ST09, GSK-ST11, and GSK-ST21) by negative selection (EasySep Human CD4+ T cell Enrichment Kit, StemCell) according to the manufacturer’s instructions. Dead cells and other debris was removed using a Dead Cell Removal Kit (Miltenyi Biotec) according to the manufacturer’s instructions. Cells from each patient were treated with 0.05% DMSO, 100nM AZD5582, or 25nM GSK445A and harvested at 2 hours, 6 hours, and 24 hours after exposure. RNA was isolated from the harvested cells using ALlPrep NDA/RNA Mini Kit (Qiagen). 200ng of RNA from each sample was checked for quality using an Agilent Bioanalyzer, with RIN scores typically >9.0 suggesting high quality RNA. These total RNA samples were then processed into stranded, mRNA libraries using the KAPA library preparation kit (KAPA BioSystems, F. Hoffmann-La Roche Ltd). Final libraries were checked by Qubit for concentration, and with a BioAnalyzer HS-DNA chip (Agilent) for fragment size distribution (mean size 359 bp). Samples were then sequenced using an Illumina Hiseq 4000 sequencer using a paired-end 50bp x 50bp run. Samples were successfully demultiplexed and then QAQCed using FASTQC (www.bioinformatics.babraham.ac.uk/projects/fastqc/).

Raw reads were mapped to the human genome and transcriptome (GRCh38.p7) using STAR and Salmon (28). Data was normalized and interrogated for changes in gene expression using DESeq2 (29) package in R. P-values were adjusted for multiple testing using a false discovery rate using the Benjamini-Hochberg method (30). Data was analyzed both jointly and within each treatment compared to the vehicle control. Differential expression of outliers was assessed and found insignificant in overall effect. Graphs and summary tables were built in R using ggplot. Gene set enrichment was performed using GSEA and GO analysis (31).

### Statistics

Statistical analysis was carried out using the pairwise, nonparametric Wilcoxon method and statistical significance was assigned to p-values of less than 0.05.

## Author Contributions

GCS and RMD designed research studies, conducted experiments, acquired data, analyzed data, and wrote the manuscript. DMI, EPB, RF, and JB designed research studies, conducted experiments, acquired data, and analyzed data. MK and CDJ acquired data and analyzed data. DF, JPR, and NMA provided reagents and contributed to experimental design and interpretation. DMM designed research studies, analyzed data, and wrote the manuscript.

## Acknowledgements

Research reported in this publication was supported primarily by Qura Therapeutics, Award Number Qura 2017_01 and 2018_01. Additional support was received from CARE, a Martin Delaney Collaboratory, the National Institute of Allergy and Infectious Diseases, National Institute of Neurological Disorders and Stroke (NINDS), National Institute on Drug Abuse (NIDA) and the National Institute of Mental Health (NIMH) of the National Institutes of Health, grant number 1UM1AI126619-01. RNA sequencing was carried out in the UNC Genomics Core Facility, supported by funds from the National Cancer Institute, grant number 5P30CA016086-42. The content is solely the responsibility of the authors and does not necessarily represent the official views of Qura Therapeutics or the National Institutes of Health. We thank Jennifer Kirchherr, Erin Stuelke, Katherine Sholtis, and Brigitte Allard for technical support. We thank Josee Girouard, Dr. Cynthia Gay, Joann Kuruc, and Caroline Baker for donor recruitment and clinical management. We thank Dr. Mary Napier and Nancie Hergert for operational support. We thank Dr Ronald Swanstrom and Dr Jill Dunham for critical reading of the manuscript. Finally, we are grateful for the contributions of the patients who have participated in these studies.

---

**Fig. S1.**
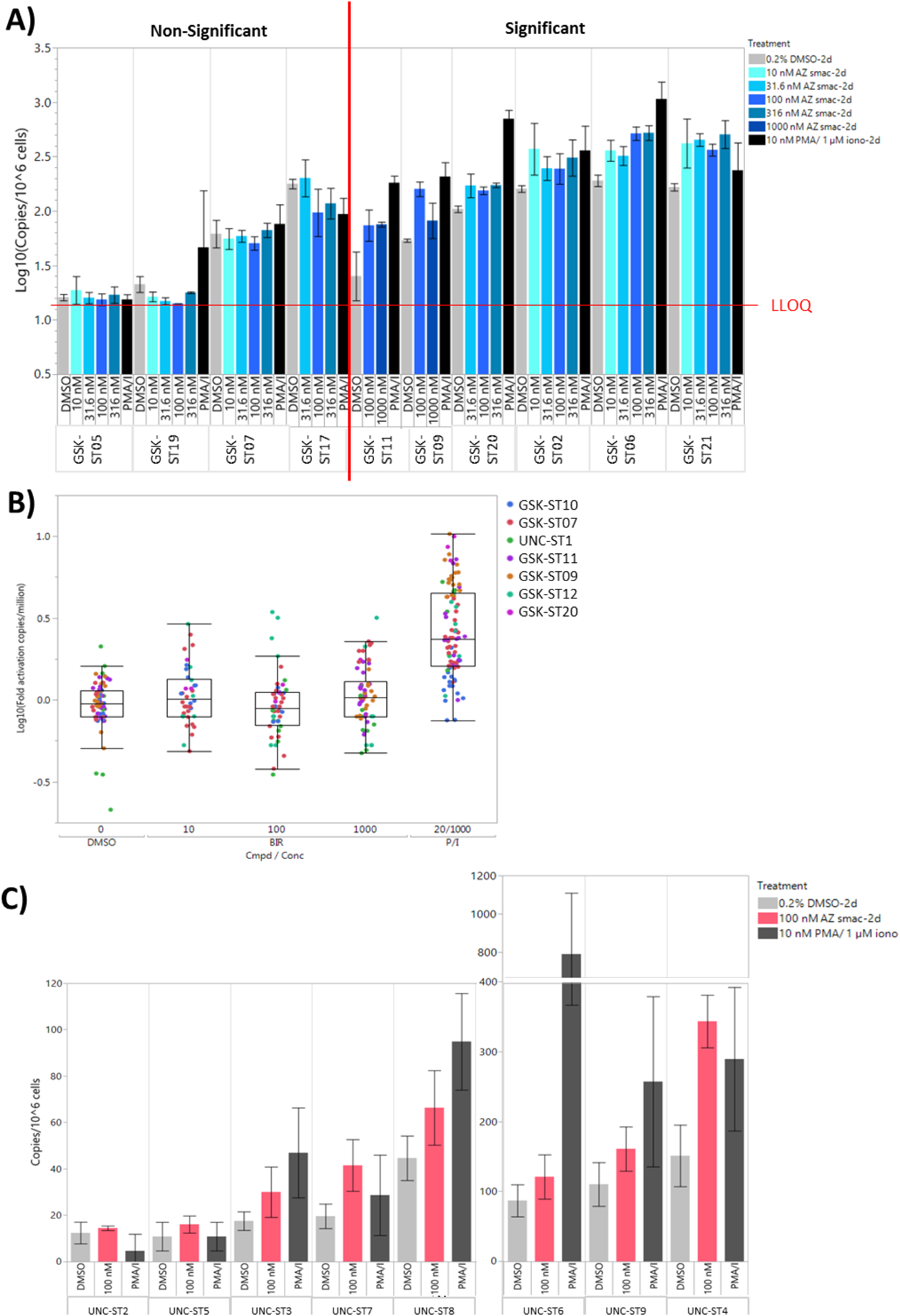
Cell associated gag RNA from total or resting CD4+ T cells treated with AZD5582 or birinapant. (A) gag RNA copies per cell in total CD4+ T cells from 10 stably treated patients treated with titrated concentrations of AZD5582 ex vivo. (B) Log-fold change in caRNA from total CD4+ T cells treated with birinapant. (C) Cell associated gag RNA copies per cell from resting CD4+ T cells of each of the eight stably-treated patients tested with AZD5582 ex vivo.

